# Imaging *Giardia intestinalis* cellular organisation using expansion microscopy revealed atypical centrin localisation

**DOI:** 10.1101/2024.01.08.574661

**Authors:** J. Soukup, M. Zelená, F. Weisz, M. Kostelanská, E. Nohýnková, P. Tůmová

## Abstract

Advanced imaging of microorganisms, including protists, is challenging due to their small size. Specimen expansion prior to imaging is thus beneficial to increase resolution and cellular details. Here, we present a sample preparation workflow for improved observations of the single-celled eukaryotic pathogen *Giardia intestinalis* (Excavata, Metamonada). The binucleated trophozoites colonize the small intestine of humans and animals and cause a diarrhoeal disease. Their remarkable morphology includes two nuclei and a pronounced microtubular cytoskeleton enabling cell motility, attachment and proliferation. By use of expansion and confocal microscopy, we resolved in a great detail subcellular structures and organelles of the parasite cell. The acquired spatial resolution of 43 nm enabled novel observations of centrin localisation at *Giardia* basal bodies. Interestingly, non-luminal centrin localization between the *Giardia* basal bodies was observed, which is an atypical eukaryotic arrangement. Our protocol includes antibody staining and can be used for the localisation of epitope-tagged proteins, as well as for differential organelle labelling by amino reactive esters. This fast and simple protocol is suitable for routine use without a superresolution microscopy equipment.

## Introduction

Expansion microscopy (ExM) is an established and commonly accessible method for improving the spatial resolution of light microscopy for biological specimens. It is based on a combination of physical and optical magnification. First described in 2015 (Chen et al. 2015), ExM has been applied to a variety of animal cells and tissues, plant cells (Hawkins et al. 2023), and was reported with several customisations for application in different research fields, such as pathology and neuroscience (Wassie et al. 2019, Ghosh et al. 2023). In general, specimens are sifted through a sodium polyacrylate mesh, the individual biomolecules crosslinked to the gel matrix and their interactions disrupted by denaturation. Then, the gel is immersed in water and the diffusion of water enables the hydrogel expansion which makes the biomolecules spread out. Labelling with fluorophores can be performed either prior to or after the expansion of the hydrogel.

ExM is a powerful approach, especially suitable for imaging microorganisms, as it can help to overcome the limitations imposed by their tiny size (Lim et al. 2019). Parasitic protists are a medically and veterinary important group of eukaryotic microbes, and ExM has been successfully applied to *Plasmodium, Trypanosoma, Leishmania, Toxoplasma* and *Trichomonas* to reveal in great detail their cytoskeletal organisation and localisation of their cytoskeleton-associated proteins (Halpern et al. 2017, Betriaux et al. 2021, Gorilák et al. 2021, Kalichava and Ochsenreiter 2021, Dos Santos Pacheco et al. 2022, Bandeira et al. 2023). Although this method has already been applied also to *Giardia* in a combination with superresolution microscopy (Halpern et al. 2018, Hardin et al. 2022), here, we present a workflow that does not require a superresolution microscopy equipment. It is based on a robust ultra-structure expansion microscopy, which is commonly used for imaging of cytoskeletal structures (Gambarotto et al. 2021) and on a modification of the method in terms of effectiveness and epitope accessibility via postexpansion labelling.

*Giardia intestinalis* (syn. *lamblia*, syn. *duodenalis*) is a binucleated parasitic protist (Metamonada, Excavata) that infects the upper intestine of humans and animals and causes a diarrhoeal disease called giardiasis. Its broad transmission range makes it one of the most frequent intestinal parasites. *Giardia* motile trophozoites possess a complex microtubule-based cytoskeleton that was imaged by different methods e.g. by light and electron microscopy in Nohýnková et al. (2006) and by ExSIM in Halpern et al. (2017). The trophozoite cell contains two nuclei (Tůmová et al. 2016) and many parasitic life-adapted organelles, such as a mesh of peripheral vacuoles (Santos et al. 2022) or highly reduced mitochondria (Tůmová et al. 2021). The ExM pipeline presented here enables imaging of the structural context of the whole *Giardia* cell, yielding high resolution information about specific protein localisation, such as that of centrin. Centrin (syn. caltractin) is a canonical basal body associated protein, which has been shown in *Giardia* to localize to two distinct clusters adjacent to the two nuclei during interphase (McInally and Dawson 2016). Here we report novel observations regarding a more precise centrin localisation in *Giardia* cells by use of expansion microscopy.

## Materials and Methods

### *Giardia intestinalis* transfection for centrin/caltractin expression

The WBc6 (ATCC 50803) cell line was used for episomal transfection and expression of the centrin (syn. caltractin) gene product (GL 50803_104685, http://giardiadb.org). The plasmid construct pTG containing HA-tagged centrin was prepared as follows: a PCR product containing the promoter (200 bp 5‘UTR) and ORF but excluding the stop codon was created using the primers 104685F (CTAGGATATCACTTTATCGTGTAACCGG) and 104685R (CTAGCTGCAGAGAGAAAGCACTTGTG) with *Eco*RV and *Pst*I sites incorporated, respectively, from *Giardia* gDNA. The PCR conditions were as follows: 98 °C for 30 s, followed by 32 cycles of 98 °C for 10 s, 56 °C for 20 s and 72 °C for 45 s, with a final extension of 72 °C for 2 minutes; Q5^®^ High-Fidelity DNA Polymerase (New England Biolabs) was used according to the manufacturer’s protocol. The PCR product of the expected size of 748 bp was purified and subcloned into the *Eco*RV/*Pst*I-digested pTG vector containing an HAHA tag at the C-terminus and a transcription unit for the selection marker puromycin-N-acetyltransferase (pac cassette) (Martincová et al., 2012). The plasmids were isolated according to the manufacturer’s instructions using a Midi Spin Column Plasmid Kit (Geneaid) and sequenced using vector-specific forward and reverse primers on a 3130xl Genetic Analyser (Applied Biosystems). Transfection of *Giardia* trophozoites was performed as follows: approx. 1 × 10^7^ trophozoites (in 300 μl of culture medium) were incubated with 50 μg of the plasmid on ice for 15 min and electroporated at 350 V, 1000 μF and 750 Ω in a 0.4 cm-gap electroporation cuvette (GenePulser Xcell Electroporation System, Bio-Rad). Transfectants were maintained under the pressure of selective antibiotics (54 μg/ml puromycin).

### Western blotting

To validate the expression of the fusion protein, cells expressing the HAHA-tagged GL 50803_104685 protein were lysed in SDS-electrophoresis buffer and proteins separated by SDS-PAGE in 12% acrylamide gel. Resolved proteins were transferred from gel onto 0.2 μm Immuno-PVDF membrane (cat. no. 162-0176, Bio-Rad) by wet western blot. The membrane was washed in Tris-buffered saline (TBS) with 0.05% (v/v) Tween-20 (TBST), saturated with 1% (w/v) BSA in TBST (TBSTB) and probed with rat monoclonal Anti-HAHA High Affinity antibody (clone 3F10, cat. no. 11867423001, Roche) diluted 1:1000 in TBSTB for 1 hour at room temperature. The membrane was washed in TBST and blocked again with BSA followed by incubation with anti-rat IgG conjugated with horseradish peroxidase (dilution 1:20000, cat. no. A5795, Sigma) in TBSTB for 1 hour at room temperature. Unbound antibody was discarded by washing the membrane in TBS. The membrane was incubated with SuperSignal West PicoPLUS Chemi Substrate (100 μl/cm^2^ of membrane, cat. no. 34579, ThermoFisher) for 5 min followed by digitization on Azure c600 (Azure Biosystems).

### Centrin localization in transfected and non-transfected *Giardia* cells

The HAHA-tagged centrin transfected and the non-transfected *Giardia* cells, as controls, were stained with rat monoclonal Anti-HAHA High Affinity antibody (clone 3F10, cat. no. 11867423001, Roche). Alternatively, to assess the endogenous central localization, both cell types were probed with mouse anti-*Chlamydomonas reinhardtii* centrin monoclonal antibody BAS 6.8 (gift from Dr. K.-F. Lechtreck, University of Köln, Germany) cross reactive to *Giardia* centrin. Used antibodies diluted in 2% BSA/PBS: Anti-HAHA tag antibody (rat) 1:250 + anti-rat AF488 conjugate (goat) 1:500; BAS 6.8 anti-centrin antibody (mouse) 1:20 + anti-mouse AF488 conjugate (goat) 1:500. Cells were imaged with Olympus IX83 microscope equipped with Hamamatsu Orca-Flash 4.0v3 16-bit camera with 100x oil immersion objective (NA 1.45). Images were taken as Z-stack with 10 μm step and Maximal Z projected with ImageJ. All images were enhanced with brightness and contrast (histogram stretching) for purpose of visualisation.

### Expansion procedure and staining strategy

The protocol was developed based on Gorilák et al. 2021 and adapted for use on *Giardia* cells. First, the trophozoites from the culture tubes (approx. 10^7^ cells) were harvested by chilling on ice for 7 min. A 12 mm coverslip (High Precision 12 mm, Marienfeld) was placed on the bottom of a 24-well plate and covered with 2 ml of the trophozoite culture. The plate was placed in a 37 °C incubator in a plastic bag with an Anaerogen sachet (Thermo Scientific™ Oxoid) to induce a microaerophilic environment. The cells were allowed to settle onto the coverslip for 30 min. Then, the cells were washed with 500 μl of PBS and fixed with 500 μl of fixative solution (4% formaldehyde, Sigma, and 4% acrylamide, Sigma, in PBS) overnight (O/N) covered in aluminium foil at room temperature (RT). Then, the cells were washed again in PBS. After removing the coverslip, the following steps were performed on parafilm in a wet chamber: gelation, denaturation and expansion of the gel were performed as described in Gorilák et al. 2022. The expanded gel (∼5 cm) was examined microscopically on an Olympus IX83 inverted microscope with a 10× or 20x phase contrast objective with a long working distance to select pieces of gel with satisfactory density of cells. An approx. 10x10 mm piece of the gel was cut out for each antibody staining. Staining was performed in a 24-well plate, and prior to staining, the gel was shrunk by incubation in 1 ml of PBS for 20 min. The volume of the staining solution was therefore reduced to 250 μl. The following mixture of primary antibodies was used: a mouse anti-acetyl-alpha tubulin antibody clone 6-11-B1 (Sigma⍰Aldrich) at a 1:100 dilution and a rabbit anti-HA tag antibody at a 1:50 dilution (Cell Signalling Technology), both in 2% BSA/PBS. The staining was performed O/N in the dark on a shaker at 200 rpm at RT. The gel was washed three times in PBS for 20 min. The secondary antibodies were applied O/N on a shaker: Alexa Fluor 594 anti-mouse IgG (H+L) and Alexa Fluor 488 anti-rabbit IgG (H+L) (both Invitrogen), both at 1:500 dilutions in 2% BSA/PBS. Three washes in ddH_2_O, each 20 min, were performed to expand to the gel to its original size. Alternatively, the pan-protein labelling reagent Atto-NHS ester 594 (Merck) was used (10 μg/ml in PBS) to non-specifically label proteins (their primary amine residues) to visualise different cell compartments and organelles. The unbound ester was washed off by three washes in ddH_2_O, each for 20 min. Negative controls of nontransfected *Giardia* trophozoites were produced. For imaging, the piece of gel was transferred to a poly-L-lysine-coated glass-bottom Petri dish (Cellvis) and imaged in a Leica TCS SP8 confocal microscope using a 63x oil immersion objective. Confocal z-stacks and individual images were acquired and processed in Fiji ImageJ software. 3D reconstructions and video display were performed in Fiji and Arivis v. 3.6 (Zeiss), and in Amira™ software (ThermoScientific). Images were enhanced with brightness and contrast (histogram stretching) for purpose of visualisation.

### Assessing the extent of expansion

To evaluate expansion extension of the ExM approach, comparative measurements of the axonemal diameter were performed on expanded and nonexpanded preparations. For the latter, the transfectant cells were fixed and stained using a standard protocol (Nohýnková et al. 2006) with the same combination of primary and secondary antibodies: mouse anti-acetyl-alpha tubulin antibody clone 6-11-B1 (Sigma⍰Aldrich) at a 1:100 dilution and rabbit anti-HA tag antibody (Cell Signalling Technology) at a 1:500 dilution in 2% BSA/PBS; Alexa Fluor 544 anti-mouse IgG (H+L) and Alexa Fluor 488 anti-rabbit IgG (H+L), both at 1:500 dilutions in 2% BSA/PBS. The preparations were mounted in Vectashield/DAPI (Vector Laboratories) and imaged with a Leica TCS SP8 confocal microscope. In total, 70 measurements for each expanded and non-expanded cell group were performed on raw images in Fiji ImageJ software. Statistical analyses were done in GraphPad Prism software v. 5.03 (GraphPad Software). Unpaired two-tailed Mann-Whitney test was used for comparison of the two datasets.

## Results

### Customising expansion microscopy for *Giardia* trophozoite cells

Previously, ExM has already been applied to *Giardia* as ExSIM, combining preexpansion labelling and structured illumination microscopy (Halper et al. 2017). Based on the approach named ultrastructure expansion microscopy (Gambarotto et al. 2021) a simpler protocol using conventional confocal microscopy and postexpansion labelled preparations has recently been introduced for unicellular kinetoplastids (Gorilák et al. 2021, Kalichava and Ochsenreiter 2021), which did not require highly specialised instrumentation, such as a superresolution microscope. Here, we adapted this protocol for use in *Giardia* research by introducing and/or modifying the following steps: (i) Instead of pelleting cells by centrifugation, the cells in their culture medium were allowed to attach to coverslips by their adhesive plates in an anaerobic environment at 37 °C. This ensured the uniform ventro/dorsal orientation of all *Giardia* trophozoites on a coverslip and prevented cell modifications by centrifugal forces. This step shortened the imaging time of the expanded cell, as it could be performed in the ventro/dorsal orientation (volume depth after expansion ∼20 μm). Imaging of cells in the anterior-posterior (∼70 μm) orientation would accordingly increase the imaging time and dataset size. (ii) After gelation and expansion, the cell density could be re-evaluated in a phase-contrast observation to select a suitable piece of gel with satisfactory cell density to be cut out for antibody staining steps. This approach helped to localise areas of the gel where no cell loss occurred during coverslip detachment from the polymerised gel. (iii) The gel shrinkage by PBS incubation before antibody staining allowed the gel to be processed in a smaller volume (24-well plate instead of a 6-well plate), which ensured that the gel was submerged completely. (iv) Although superresolution microscopy was not applied, imaging with a conventional confocal microscope yielded satisfactory results for our applications, allowing to discern individual microtubules (unexpanded size: Ø25 nm) of the ventral disk, funis and flagellar axonemes.

### Expansion microscopy resolved *Giardia* cellular structures

The ExM method relies mostly on antibody staining; in our experiments, the anti-acetylated tubulin (6-11-B-1 Ab) antibody was used to decorate *Giardia* microtubular cytoskeleton. This antibody recognizes all microtubular systems in *Giardia*, including eight basal body/flagellum, ventral disk, median body, funis and mitotic spindle microtubules (Marková et al. 2016) (**Fig. 1a**). Additionally, the protein labelling reagent Atto-NHS ester 594 was applied to our samples to visualise different organelles, such as the two nuclei, peripheral vacuolar system and the cytoskeleton (**Fig. 1b**). Our protocol provided a substantial improvement in light-microscopy resolution of the *Giardia* cellular structures. 3D rendering of the data and ventral- and dorsal-views with different opacities are shown (**Fig. 1c, d**). The entire volume of *Giardia* trophozoite cells was imaged at relatively high resolution (pixel size: xy 43 nm and voxel depth: 300 nm). The cells increased their volume and organelle size by a factor of 3.4 (p-value < 0.0001) according to measurements of axonemal diameters (**Suppl. Fig. 1**). Comparison of expanded and nonexpanded cells illustrated in **Fig. 2** and **Suppl. Video 1 and 2** demonstrated cellular details under the same magnification (e.g. the individual microtubular doublets forming an axoneme, microtubules of the adhesive plate and funis, or centrin localization can be visualized with a substantial better resolution in expanded cells).

**Fig 1.**
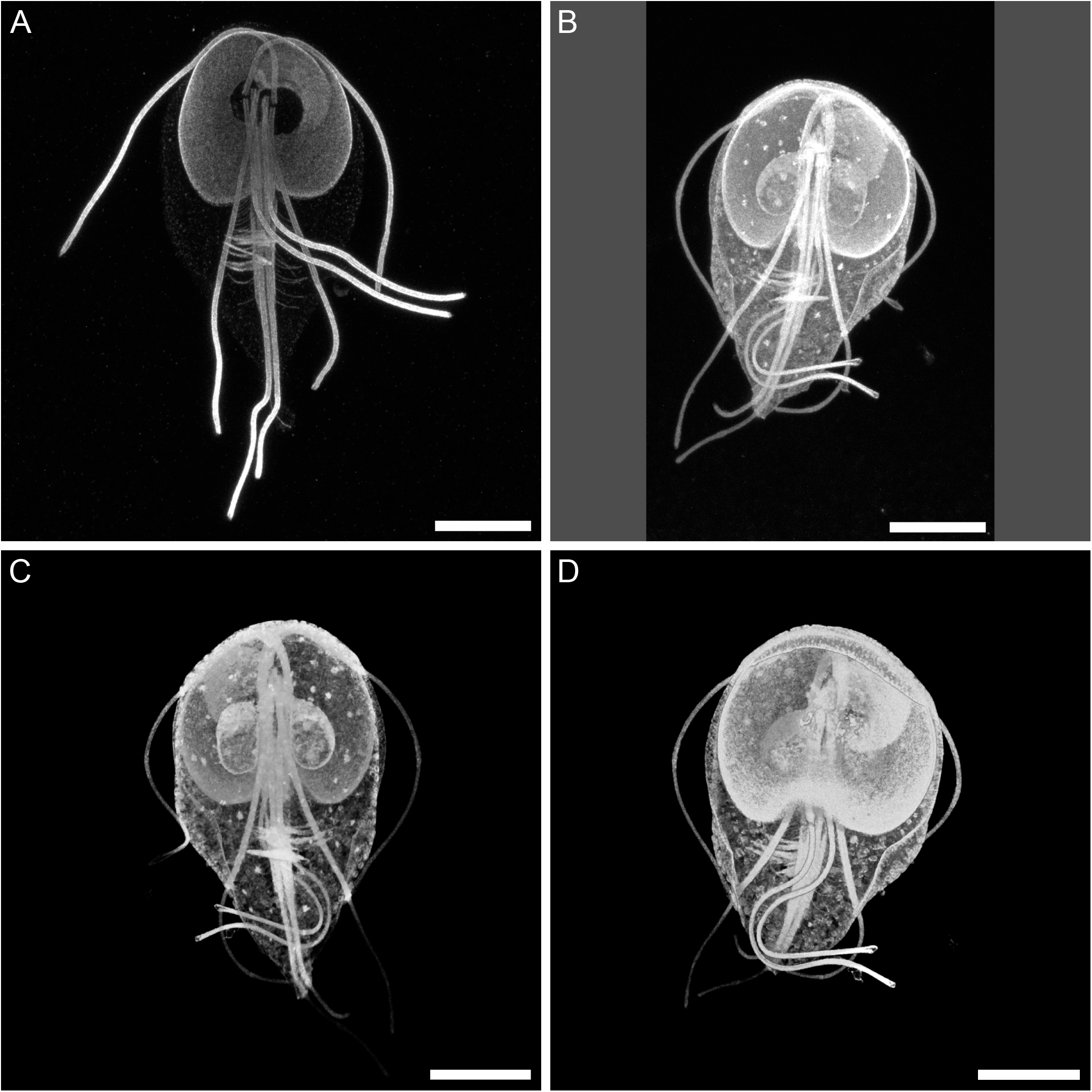
Resolving *Giardia intestinalis* cellular structures by ExM. A: Maximal projection of confocal z stack of antibody labelled microtubules (antibody 6-11-B1 against acetylated tubulin) of expanded *Giardia* trophozoite, B: Maximal Z projection of confocal images of an expanded *Giardia* trophozoite labelled by NHS ester binding primary amines. Scale bars: 20 μm. Image B was underlaid with a grey square the original image is shown). C and D: 3D reconstruction of NHS ester labelled and an expanded *Giardia* cell showing its dorsal (C) and ventral (D) side. 3D reconstructions and volume rendering were done in Amira software (ThermoFisher).

**Fig 2.**
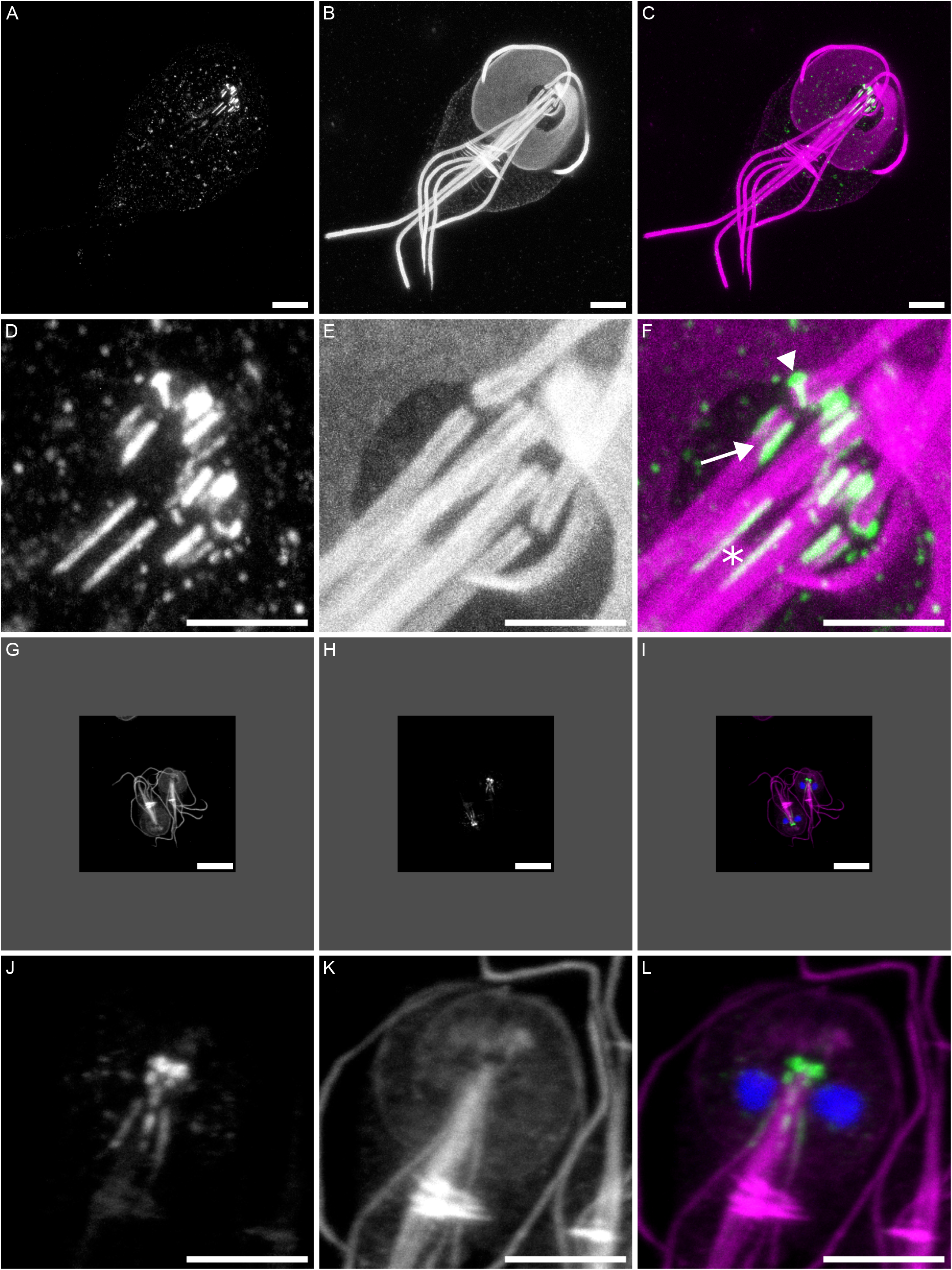
Comparison of centrin localization in expanded and non-expanded *Giardia intestinalis* cells. Example images comparing ExM images of *Giardia* trophozoites transfected with centrin-HAHA and standard immunofluorescent images of non-expanded cells obtained by a widefield microscope. A: An image of an ExM cell labelled with anti-HAHA. B: Identical cell as in A, labelled with anti-acetylated tubulin antibody 6-11-B1. C: Merged image of A and B (green - centrin, magenta - tubulin). D - F: detail of the area containing centrin in the cell shown in A-C. One longer and one shorter fibre longitudinally accompany the microtubular triplets of all basal bodies (arrow), except those of anterolateral flagella. There, a peri-basal body capping or ring is present, ending as an accumulation of centrin material directed toward the nucleus (arrowhead). Two fibres run along the upper part of caudal axonemes (asterix). G - I: Standard immunofluorescence of non-ExM cell with labelled centrin (anti-HAHA antibody) (G), acetylated tubulin (H) and merged image with DAPI (I). J-L - detail of centrin area of *Giardia* cell shown in G-I. Despite the same magnification of images C and I, and of images F and L, the non-ExM cell (L) does not provide as many details on centrin localization as the ExM cell (F). Figure G-I - the original images are shown in order to match scale bar size with the cell in A - C. The scalebars: A-C and G-I - bars 10 μm. D-F and J-L - bars 5 μm.

**Fig. 3.**
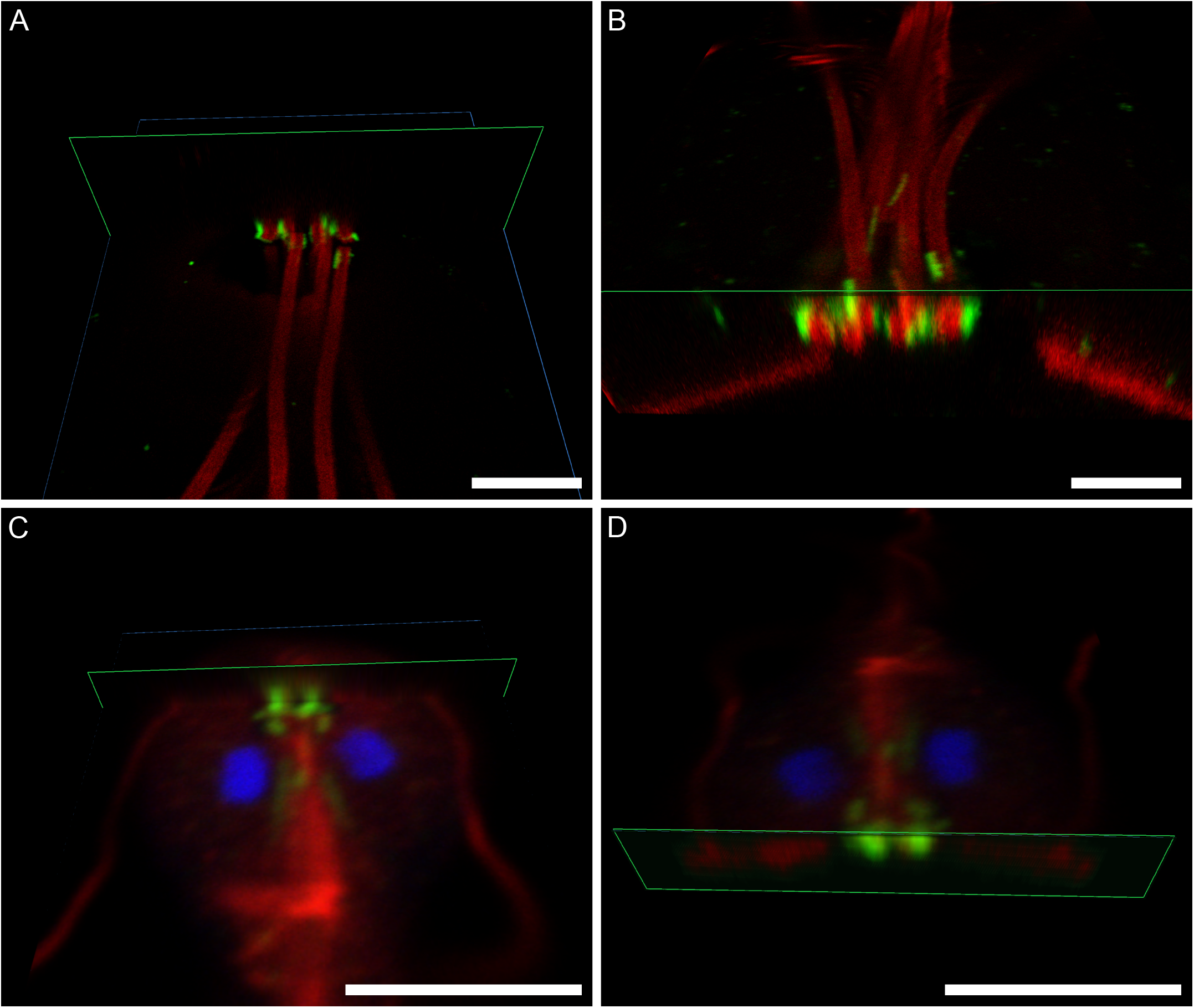
3D reconstruction of centrin localization in expanded and non-expanded *Giardia intestinalis* cells. Example images from 3D rendering from Arivis software. A and B: ExM confocal z-stack 3D rendering with XY (blue) and XZ (green) planes (A) and cropped 3D reconstruction of a cell (B). C and D: confocal z-stack of a non-expanded cell 3D rendering with XY and XZ planes (C) and cropped 3D reconstruction of cell (D). Scale bar 5 μm.

### Centrin localisation in *Giardia* demonstrated by ExM

As an example of usefulness of our ExM approach a more precise localisation of *Giardia* centrin was determined, which is a substantial improvement over conventional wide-field or confocal microscopy centrin imaging in non-expanded cells (**Fig. 2, Suppl. Fig. 3, Suppl. Video 1 and 2**). Centrin (GL 50803_104685, syn. caltractin) was selected based on its stable interaction with microtubule organizing centres (MTOCs), such as basal bodies in *Giardia*, and expressed as epitope-tagged protein under its native promoter. Expression of the fusion protein in transfectant cells was confirmed on Western blots as 22 kDa protein (**Suppl. Fig. 2**), and its intracellular localization by widefield and confocal microscopy (**Fig. 2, Suppl. Fig. 3**). Additionally, staining with anti-*Chlamydomonas* centrin antibody (BAS 6.8), previously shown to cross react with *Giardia* centrin, was applied to transfectants and non-transfected cells to verify the centrin localization in trophozoite cells (**Suppl. Fig. 4**).

By use of ExM, we found that *Giardia* centrin is localised between the basal bodies of the eight flagella, at the anterior part of the trophozoite cell. One longer and one shorter fibre stretch alongside the outer surface of the microtubular triplets of all basal bodies except those of the anterolateral flagella. There, a peri-basal body capping or ring is present, ending as a stronger accumulation of centrin material directed toward the nucleus. Additional centrin accumulation is present also between the distal part of the anterolateral basal body and a proximal part of the caudal basal body. Two tiny fibres run along the upper part of the caudal axonemes. However, no connection to the centrin at basal bodies was observed (**Fig. 2d**). As confirmed in 3D-projections, no staining inside the basal bodies was found, which indicates a non-luminal localization of *Giardia* centrin (**Fig. 3, Suppl. Video 1**).

## Discussion

Expansion microscopy achieves via a specific sample preparation procedure an increase resolution of cellular structures; we demonstrated applicability of its use for *Giardia intestinalis* cells based on the modification of the ultrastructure expansion microscopy protocol previously successfully applied to other protists. Physical expansion combined with post-expansion labelling and confocal microscopy increased the resolution of resulting images by the factor of 3.4, which is slightly less than the usual expansion index of 4.5 (Wassie et al. 2019), however we demonstrate that it clearly enables resolving of details previously not accessible by conventional light microscopy. The post-expansion labelling generally improves the epitope accessibility of expanded samples and reduces antibody linkage errors proportionally to the expansion factor (Sauer 2021), and was therefore preferred in our experiments over pre-expansion labelling previously used in *Giardia* (Hardin et al. 2019, 2022). Our protocol utilizes a conventional confocal microscope routinely present in microscopy facilities instead of less common superresolution devices. Antibodies and an amine-reactive dye were shown to label the *Giardia* subcellular compartments and organelles. The achieved resolution after expansion enabled discerning individual microtubules, microtubule doublets of flagellar axonemes and specifically, the precise localisation of centrin in *Giardia* cells expressing a tagged variant of the protein.

Centrins are ubiquitous, highly conserved calcium-binding phosphoproteins associated with MTOCs, such as the centrosome of non-ciliated cells, the basal bodies of ciliated or flagellated cells, and the spindle pole body of yeast cells (Salisbury et al. 1988, Moretti et al. 2023). In mammalian cells, centrin is present in the lumen of the centriole in its central core (Le Guennec et al. 2020). In algal cells, centrins are localised mostly as nucleus–basal body connectors, distal striated fibres and at flagellar transition regions (Taillon et al. 1992). By use of immunoelectron microscopy, a continuous filamentous scaffold that extends from the nucleus to the base of axonemes was described in *Chlamydomonas*. In this alga, also a rotational asymmetry of centrin distribution in the distal lumen of the basal bodies was described (Geimer and Malconian 2005).

Based on its localization in the trophozoite cell, centrin in *Giardia* was for many years hypothesised to play a role in motility and basal body positioning. *Giardia* has two centrins functions of which are not completely understood. Human and algal centrin antibodies immunolocalised *Giardia* centrin to the basal bodies, posterolateral axonemes and in some studies to the median body (Belhadri 1995, Meng et al. 1996, Correa et al. 2004). Since heterologous antibodies recognised both centrins, Lauwaet et al. (2011) individually epitope tagged both centrins and found that centrins 1 (Gene ID GL50803_ 6744) and 2 (Gene ID GL50803_ 104685) have identical localisation patterns and localise to the basal bodies and posterolateral axonemes. On the contrary, in GFP centrin transfectants (GL 50803_104685/GFP) (McInally and Dawson, 2016), no staining was shown alongside the posterolateral axonemes. In the above-mentioned works, the resolution of light microscopy was unable to provide a detailed assessment of centrin localisation regarding the basal bodies. In our study, the detected centrin localisation by expansion microscopy corresponds to that described by McInally and Dawson (2016) for GL 50803_104685/GFP transfectants, i.e. without the signal alongside the posterolateral flagellar axonemes. These epitopes were probably lost in expansion experiments by lowering their concentration during expansion. By comparing localization of overexpressed and endogenous centrin in *Giardia* by widefield microscopy, it could be determined that at the posterolateral axonemes there is some storage or accumulation of centrin. *Giardia* trophozoites possess in their eight flagella different paraflagellar structures, e.g. dense rods accompanying the intracytoplasmic portions of the posterolateral flagella (Kulda and Nohýnková 1995). Many proteins localizing to these structures were identified, including intraflagellar transport proteins (Hagen et al. 2020). We found short centrin fibres also alongside the caudal axonemes of *Giardia* trophozoites. To what extend it helps the *Giardia* cytoskeleton stabilization or signalling, is not clear.

Regarding the basal bodies-associated centrin localization in *Giardia*, an interesting observation resulted from our ExM work. We found centrin linkers among all basal bodies and around basal bodies of the anterolateral flagella. Linkers exist also between anterolateral basal bodies and *Giardia* nuclei. In contrast to mammalian and algae cells (Bornens and Azimzadech, 2007, Laporte et al. 2022), the 3D reconstructions of expanded *Giardia* basal bodies did not show any luminal centrin localization inside the basal bodies. A circular scaffold inside the centrioles or basal bodies, which involved centrin in the central region, was shown to resist deformation forces to help hold microtubule triplets together in the mammalian and algae centrioles/basal bodies (Le Guennec et al. 2020). In *Giardia*, no information is available on a precise localization of proteins to the basal body inner scaffold or specifically, to proximal, central or distal regions of the eight basal bodies of a trophozoite cell.

The exact purpose of the described organisation is unclear, but it is possible that centrin provides a mechanical support for flagellar beating, interconnects and stabilizes the mastigont (all the axonemes) and provides a link to the *Giardia* nuclei. Further functional studies will be needed to investigate its hypothesized role in *Giardia* movement and flagellar apparatus stabilisation.

## Acknowledgements

The authors declare no conflict of interest. We acknowledge Eva Pyrihová for support with transfection experiments, Vladimír Varga for critical reading of the manuscript, and the Laboratory of Confocal and Fluorescence Microscopy, Faculty of Science, CUNI. The work was financially supported by Czech Science Foundation (GAČR) grant 20-06498S to E.N., by PRIMUS/20/MED/08 to P.T. and by Charles University Research Centre program No. UNCE/24/SCI/011.

## Figure Legends

<u>Supplementary Fig. 1. Assessing of the extend of expansion of flagellar axoneme diameters.</u>

Multiple positions alongside the axoneme lengths were measured. 70 measurements were done in total in 8 expanded cells and 10 non-expanded cells. Statistical analysis was done in GraphPad Prism software v. 5.03 (GraphPad Software). Unpaired two-tailed Mann-Whitney test was used for comparison. Each spot represents 1 measurement. The thick line represent mean with standard deviation. *** p-value < 0.0001.

<u>Supplementary Fig. 2. Detection of the tagged centrin in transfectant cells on a Western blot.</u>

Image of Western blot probed with a rat monoclonal anti-HAHA antibody and developed with anti-rat IgG conjugated with horseradish peroxidase on *Giardia* cells (transfected WBc6 cell line) expressing the HAHA-tagged centrin. The 22 kDa centrin band is present. Marker: PageRuler™ Prestained Protein Ladder, 10 to 180 kDa.

<u>Supplementary Fig. 3. Details of a centrin localization in expanded</u>*<u>Giardia</u>* <u>cells.</u>

Cropped details of the area containing centrin in an expanded *Giardia* trophozoites cell expressing a centrin-HAHA. Green: anti-HAHA antibody. Magenta: anti-acetylated tubulin antibody. One longer and one shorter fibre longitudinally accompany the microtubular triplets of all basal bodies (arrow), except those of anterolateral flagella. There, a peri-basal body capping or ring is present, ending as an accumulation of centrin material directed toward the nucleus (arrowhead). Two fibres run along the upper part of caudal axonemes (asterix).

Bars 10 μm.

<u>Supplementary Fig. 4 Centrin localization in non-expanded</u>*<u>Giardia</u>* <u>cells.</u>

Example widefield images of HAHA-tagged centrin expressing *Giardia* cells stained with anti-HAHA antibody (A-C) and with BAS 6.8 anti-*Chlamydomona*s centrin antibody (cross reactive with *Giardia* centri*n*) (D-F). Used antibodies: Anti-HAHA tag antibody (rat) + anti-rat AF488 conjugate (goat); BAS 6.8 anti-centrin antibody (mouse) + anti-mouse AF488 conjugate (goat). Bars: 10 μm. Cells were imaged with Olympus IX83 microscope equipped with Hamamatsu Orca-Flash 4.0v3 16-bit camera under 100x oil immersion objective (NA 1.45). Images were taken as Z-stack with a 10 μm step and Maximal Z projected with ImageJ.

