## Supplementary figures and images for "Imaging *Giardia intestinalis* cellular organisation using expansion microscopy revealed atypical centrin localisation"

### Supplementary Fig. 1

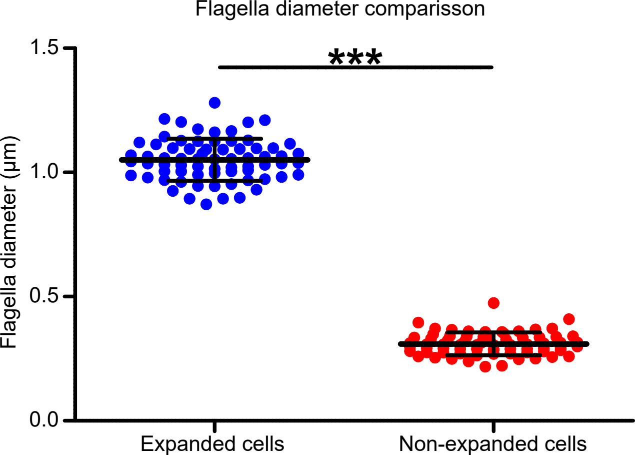

### Supplementary Fig. 2

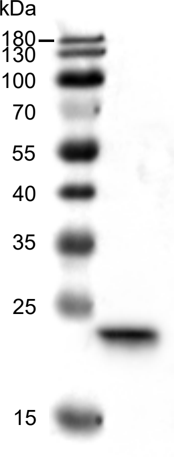

### Supplementary Fig. 3

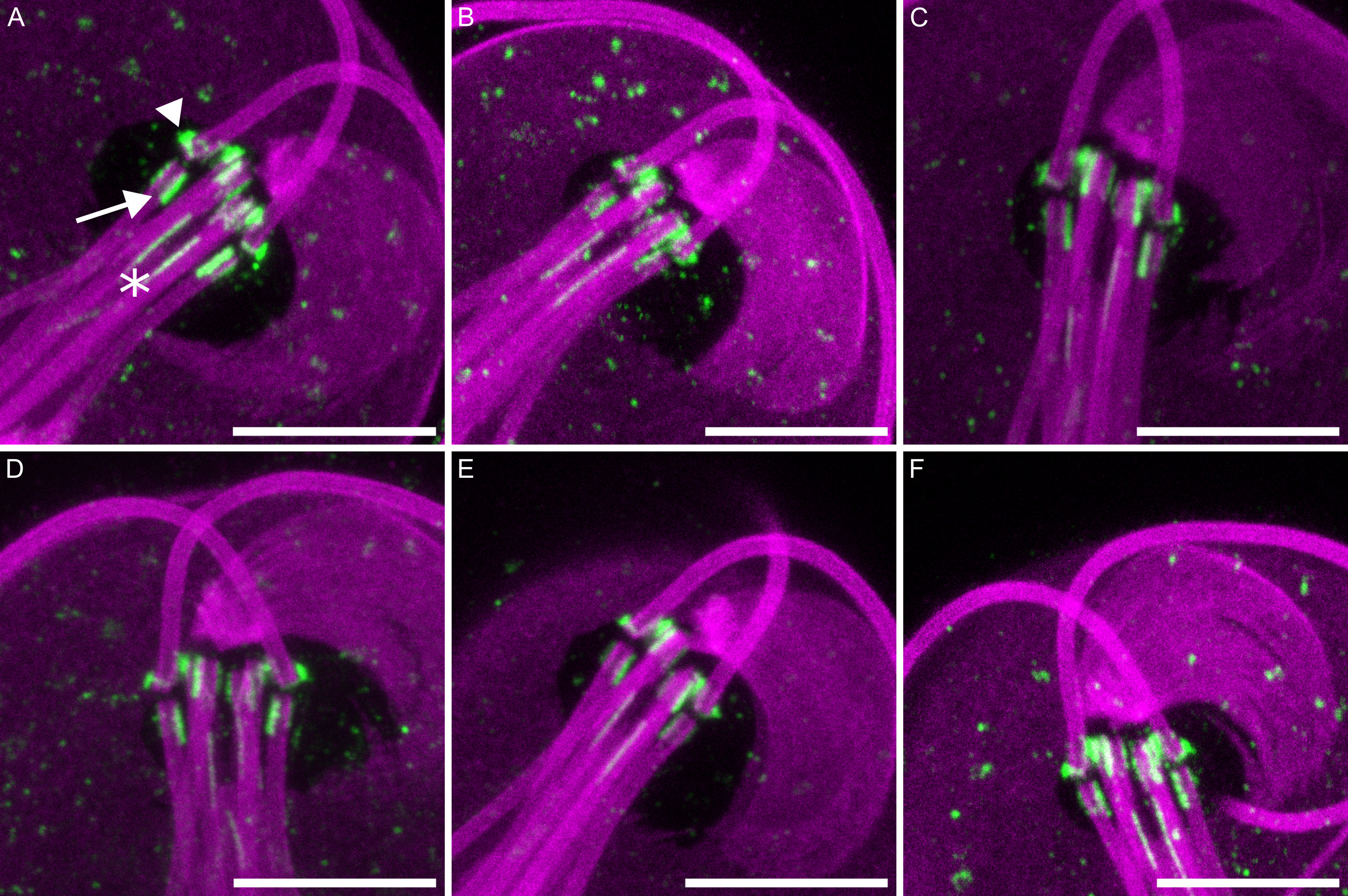

### Supplementary Fig. 4

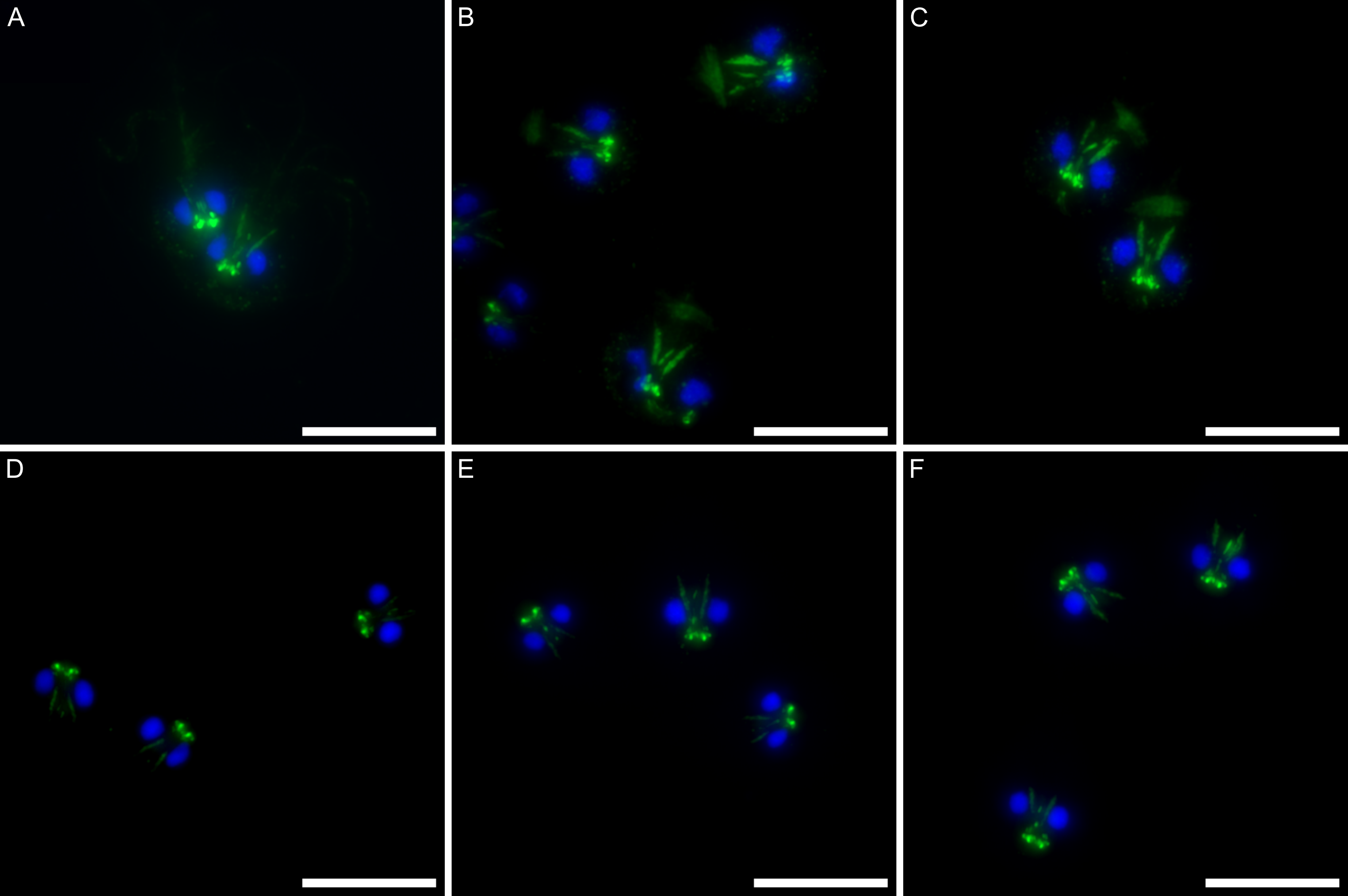
